# Designing Novel and Potent Inhibitors for Multi Drug-Resistant Tuberculosis: A Computational Approach

**DOI:** 10.1101/2021.05.05.442727

**Authors:** Muhammad Idrees, Bashir Ahmad, Muhammad Israr, Muhammad Waqas, Syed Muhammad Mukarram Shah, Aishma Khattak, Saad Ahmad Khan

**Author notes:** Corresponding author Dr. Muhammad Israr, Associate Professor, Department of Biology, The University of Haripur, Khyber Pakhtunkhwa Pakistan.

## Abstract

Tuberculosis is a major global health problem and is still among the top 10 causes of death. The increasing rate of drug resistance to infectious agents has provoked an urgent need to discover novel anti-tuberculosis agents with novel modes of action. In this study, small molecule inhibitors of the proteins encoded by the drug resistant genes, i.e., katG, gyrA, pncA and rpoB of Mycobacterium tuberculosis (M. tuberculosis), were identified using computational methods. In the ligand base pharmacophore, an already reported four ligands for the four proteins encoded by the resistant genes of M. tuberculosis were selected for the generation of pharmacophores. The validated pharmacophores model of all the four proteins, generated on the basis of ligand base, were selected for further screening of ZINC drug like database. As screening results 486 structurally diverse hits for katG, 542 for PncA, 112 for rpoB and 365 for gyrA were mapped and filtered via Lipinski’s rule of five. Finally, on the basis of docking score and binding interactions, ten small molecules were selected for each protein as novel inhibitors. These selected novel inhibitors have significant interaction with the active site of the protein and a strong possibility to act as an additional starting opinion in the development of new and potential inhibitors. The result indicates that novel inhibitors could be a promising lead compound and be effective in treating sensitive as well as multi drug resistant tuberculosis.

## Introduction

Tuberculosis is a major global health concern and can be transmitted through droplet aerosol from person to person (Khan et al., 2013). The causative agent of the disastrous tuberculosis diseases Mycobacterium tuberculosis complex (MTBC). Mycobacterium bovis is another mycobacterium that can cause TB disease in people. One of the studies performed to conclude the prevalence and associated risk factors of bovine TB caused by M. bovis in most populated and centrally located districts of Khyber Pakhtunkhwa (KP) in Pakistan (AsadUllahet al. 2019).

Multidrug resistant TB develops due to mutations in the genes or a change in the titration of the drug (Migliori et al., 2007). The resistant strains are known as multiple drug resistance tuberculosis (MDR-TB) strains when they develop resistance to at least two of the first line treatment (isoniazid and rifampicin) (Prasad, 2005). When in addition to MDR, M. tuberculosis strain develops resistance to at least one fluoroquinolone and at least one of the three second-line injectable drugs, viz., amikacin, kanamycin, and capreomycin, is called extensively drug-resistant tuberculosis (XDR-TB)(Hamilton et al., 2007).

Isoniazid resistance is a complex process and develops as a result of mutations in inhA, katG, kasA, ahpC and ndh genes (Da Silva and Palomino, 2011). katG codes for the enzyme catalase/peroxidase, which is involved in the activation of isoniazid. The reduced or absent activity of this enzyme occurs due to the mutations in katG gene, which is responsible for isoniazid resistance (Zhang et al., 1992). Rifampicin works by binding with RNA polymerase β-subunit encoded by rpoBgene and thus blocking the elongation of messenger RNA (Blanchard, 1996). Pyrazinamide is equivalent to nicotinamide in terms of structure and hence naturally gets converted into an active form, pyrazinoic acid (POA) by pyrazinamidase/nicotinamidase that is encoded by pncA gene. Mutations in pncA gene leads to the loss of PZase activity, thus inhibiting the conversion of PZA to POA (Cheng et al., 2000; Juréen et al., 2008). Ofloxacin, a synthetic antibiotic of the fluoroquinolone drug class, is the second line drug that works by binding to gyrase-DNA complexes and inhibiting DNA replication. Ofloxacin resistant strains develop through mutations in quinolone resistance-determining regions of gyrA and gyrB genes (Maruri et al., 2012).

Identifying the disease causing target protein is very critical in an in silico drug designing method. The method used for investigating the protein ligand interactions is molecular docking. The insilico drug designing approach has been getting recognition as an important tool to identify potential novel drugs for various diseases (Gore and Desai, 2014). Computer Aided Drug Designing (CADD) has effectively been used in molecular biology, nanotechnology, and biochemistry and sooner it will be a dominant and commonly used technique in modern medical sciences.

The in-silico screening is also known as virtual screening (VS). It exploits computer aided approaches to identify new ligands based on biological structures. The virtual screening is either a docking method based on structural interactions or a ligand based screening. The ligand based virtual screening techniques are based on comparing the molecular similarity of compounds with known and unknown moiety irrespective of the used algorithm. Generally, the ligand-based virtual screening employs searching molecules with structural similarity to known molecules fitting well into the target binding site and hence increasing the chances of binding the target. In the present study, an effort was made to find out powerful novel small molecule inhibitors of the proteins encoded by drug resistant genes using an in-silico approach.

## Materials and Methods

Computer aided drug designing tools were applied to find out the small molecule inhibitors of the proteins encoded by drug resistant genes, i.e., katG, gyrA, pncA and rpoBof M. tuberculosis. An HP Z620 Workstations with NVIDIA DDR5, 4GB GTX–980 graphics card was used on Microsoft Windows 10 operating system. Molecular Operating Environment (MOE) software was used for screening and molecular docking. MOE was used for constructing protein and side-chain refinement. The mutants of katG, gyrA, pncA and rpoB gene of M. tuberculosis reported previously were selected for designing novel small drug inhibitors(Ahmad et al.; Ahmad et al., 2017).

### Proteins Crystal Structure Retrieving

Crystal structure present in the RCSB PDB server was used as the basis for selecting all the four proteins of M. tuberculosis. The catalase-peroxidase protein encoded by katG gene is associated with isoniazid resistance. The crystal structure of the catalase-peroxidase protein was obtained from the RCSB PDB server with PDB ID 2CCA at a resolution of 2.0 angstrom.

As follow the sequencing data reported previously, the mutations of katG gene as follows Ser303Trp and Lys274Argin variant 1, Ser315Arg in variant 2, Ser315Thr mutation in variant 3 and Gly316Ser in variant 4(Ahmad et al., 2017). The mutated structures of the four variants were observed to be created from 2CCA. In the first mutated structure, the observed alterations were Lys274Arg and Ser303Trp in protein builder of MOE and the mutated residues were found to be reduced. In the second mutated structure from 2CCA, the changes were Ser315Arg in protein builder of MOE and the mutated residue was reduced. In the third mutated structure from 2CCA, the change was Ser315Thr in the protein builder of MOE and the THR was minimized. In final mutated structure from 2CCA, the change was Gly316Ser in protein builder of MOE. After auto correction, all the structures were aligned and superposed. The pair wise RMSD plot was created using MOE.

pncAgene codes for the enzyme associated with pyrazinamide drug processing, nicotinamidase/pyrazinamidase. The crystal structure of nicotinamidase/pyrazinamidase was obtained from the RCSB PDB server with PDB ID 3PL1 at a resolution of 2.2 angstrom. As follow the sequencing data reported previously, the reported mutations were: Ser65Ser and Cys138stop mutation in variant 1, double mutation of Ser65Ser* and Gly132Ser in variant 2, insertion of G at position 131 positions in variant 3, a Gln141Pro mutation in variant 4(Ahmad et al.). The structures of two mutated variants were created from the crystal structure 3PL1. The protein builder in MOE2016 was applied to generate mutations. The first structural mutation generated was the insertion of G at position 131 and the mutated residue was reduced. The second structural mutation generated was Gly132Ser and Gln141Pro and the mutated residues were reduced. MOE was used to autocorrect both the mutated structures by eliminating the bad interactions. 3PL1 in MOE2016 was used to align and superimpose the mutated structures and generate the pairwise RMSD plot.

The rpoB gene codes for a protein named DNA-directed RNA polymerase β-subunit, which is linked to rifampicin resistance. The crystal structure of RNA polymerase β-subunit was obtained from RCSB PDB with PDB ID 5UHC at a resolution of 3.8 angstrom.

As follow the sequencing data reported previously, the reported mutations were: Ser450Gln in variant 1, Ser450Leu in variant 2, double mutation of Ser450Leu and Pro454His in variant 3, double mutation of Ser450Leu and Gly455Asp in variant 4 and Asp435Gly in variant 5(Ahmad et al.). 5UHC with protein builder of MOE 2016 was used to generate all mutated structures and the mutated residues were reduced. All five mutated structures were autocorrected in MOE to get rid of any bad interaction. The PDB ID 5UHC was used to align and superimpose crystal structure with the five mutated structures and generate a pair wise RMSD plot in MOE.

The gyrA gene codes for DNA gyrase subunit A protein that is linked with ofloxacin (fluoroquinolones) resistance. The gyrAprotein crystal structure was obtained from RCSB PDB server with PDB ID 3IFZ at a resolution of 2.7 angstrom. The sequencing results of gyrA gene reported previously were: Asp94Gly and Ser95Thr mutation(Ahmad et al., 2017). The 3IFZ structural mutations were found to be at 94 and 95 amino acid positions by protein builder in MOE and the mutated residues were minimized. The auto correction was applied to the mutated structure to eliminate any bad interactions. The crystal structure 3IFZ was aligned and superimposed with the mutated structure and a pair wise RMSD plot was created.

The missing loops in all structures were constructed by the loop modeling in MOE 2016 with default parameters and Amber99 force field was applied for loop generation. All the structures were auto-corrected in MOE to get rid of any bad contact or missing bonds.

### Ligand Base Pharmacophore Generation and Validation

Pharmacophore is a group of steric and electronic features that is required for the action of the drug (Khedkar et al., 2007). In the ligand base pharmacophore, an already reported four ligands for the four proteins coded by the resistant genes of M. tuberculosis were selected for the generation of pharmacophores. To evaluate the quality of a pharmacophore model it must be validated. Two methods were used for the validation of the pharmacophoric model. First, a test database of active ligands reported for the protein and non-active/ least active compounds were scanned on the pharmacophoric features of the ligands. In the second procedure of ligand validation, the presence of the important chemical features present in the pharmacophore which interact with important amino acids in the active pocket of the corresponding receptor protein was also used for the validation of the pharmacophore model (Wadood et al., 2017). Isoniazid is a reported ligand for katG protein. The selected pharmacophoric features were three hydrogen bond acceptors, two hydrogen bond donors, one aromatic ring interaction and one hydrophobic interaction by MOE 2016 following the default protocol. The seven characteristics for katG were selected as essential. A test database of 12 reported katG inhibitors were used for validation of generated model (Vilchèze et al., 2011). The test database containing 12 active and 12 non-active/ least active compounds were tested on the 7 featured pharmacophore and evaluated their mapping modes. In the important features of ligand pharmacophore validation, the pharmacophoric features were selected on the bases of interaction with the protein residues present in the crystal structure of the katG protein. The reported ligand for pncA protein is pyrazinamide. There are seven pharmacophoric characteristics present in pyrazinamide: three hydrogen bond acceptors, one hydrogen bond donor, two hydrophobic interactions and one aromatic interaction by MOE2016 following the default parameters. All the characteristics for the pharmacophore of pncA gene were considered to be essential. For the validation of the pharmacophore, a test database was selected with six active ligands and six non-active/ least active compounds and the pharmacophore was screen on the test database (Seiner et al., 2010). In the important features of pyrazinamide ligand pharmacophore validation, the pharmacophoric features were selected on the bases of interaction with the protein residues present in the crystal structure of the pncA protein. Rifampicin is the reported ligand for rpoB protein. The pharmacophoric characteristics of rifampicin ligand contain three hydrogen bond acceptors, three hydrogen bond donors, one aromatic interaction and one hydrophobic interaction by MOE2016 following the default protocols. In eight pharmacophoric characteristics five were selected as essential. For the validation of the ligand model generated, all the features were selected those which interact with the important residues of the active site residues atoms in the crystal structure of rpoB protein. Ofloxacin (a fluoroquinolone) is the reported ligand for gyrA protein. The pharmacophoric characteristics of ofloxacin are three hydrogen bond acceptors, one hydrogen bond donor, two aromatic interactions and one hydrophobic interaction by MOE2016 following the default parameter. All features were selected as essential for the pharmacophore. For the validation of the pharmacophore 22 known inhibitors were selected in a test database and screened on the pharmacophore. The 11 actives and 11 non-active compounds were selected for screening and all the 11 active compounds were selected by the pharmacophore modeled (Anderle et al., 2008). The features selected for the pharmacophore were on the bases of the ligand atom interaction with the active site residues atoms.

### Complexed base Pharmacophore Screening

The pharmacophores of all the four proteins, generated on the basis of ligand base, were selected for further screening of ZINC small molecule drug-like a database comprises of 17,900,742 small molecules. To confirm whether the hits retrieved from the database retain drug-like properties, all the obtained hits in all the four output databases were scrutinized for Lipinski’s rules of five. All these pharmacophores were chosen for additional evaluation.

### Molecular Docking

The retrieved outputs from the pharmacophore databases were docked in the proteins active site. The katG encoded protein has the active site corresponding to the UniProt server accession number P9WIE5. The residue positions 104, 108 and 321 are from the active site while 270 is a metal binding site. The crystal structure of the katG protein with PDB ID 2CCA shows the ligand bonded in the active site. These ligands bonded residue pockets were further evaluated for docking of katG output screen database. The docked hits retrieved from katG output databases were saved in a new database. For each ligand, five conformations were saved applying the proxy triangle algorithm that followed the London dG scoring methodology for refinement in MOE2016.

The crystal structure of pncA protein corresponding to UniProt accession number I6XD65 has the active site residues, at 8, 96 and 138, while the metal-binding residues are positioned at 49, 51, 57, and 71. The crystal structure with PDB ID 3PL1 was evaluated for docking the pharmacophore output database. The residues at 8, 96, and 138 were selected as an active pocket. The output database was docked in the active pocket and the results were saved in a new output database. For each ligand, five conformations were generated using the proxy triangle algorithm, followed by London dG scoring methodology for refinement in MOE2016.

The DNA-directed RNA polymerase β-subunit protein with PDB ID 5UHC contains 429, 435, 438, 439, 451, 454 and 465 residues at the active pocket. The ligand rifampicin was docked in the active pocket of crystal structure corresponding to 5HUC. The active site residues obtained from the ligand base screening were selected for docking, and a fresh output database was generated from the docked hits. For each ligand, five conformations were selected using the proxy triangle algorithm, followed by London dG scoring methodology for refinement in MOE2016.

In the DNA gyrase subunit A protein, the active site amino acid residue is positioned at 129. For docking, a crystal structure (PDB ID 3IFZ) was selected. The new library was created from the docking outputs. The active site amino acid residue (129) was selected for docking with the output database. For each ligand, five conformations were selected using the proxy triangle algorithm, followed by London dG scoring methodology for refinement in MOE2016.

### Binding Affinity and Energy Calculations

The binding affinities were calculated using the generalized Born/volume integral (GB/VI) implicit solvent algorithm implemented in the MOE2016. Generally, the non-bonded interaction energy between ligand molecules and the protein residues is referred to as the generalized Born interaction energy, which comes from the Van der Waals forces, Coulomb interaction, and implicit solvent interaction energies. However, protein residues and ligand strain energy are not ever taken into the calculations. After energy minimization, the binding affinity was calculated for each hit and stated in the unit of Kcal/Mol.

## Results

### Crystal Structures of M. Tuberculosis Resistant Gene’s proteins

The four different mutated structures were acquired from the crystal structure 2CCA (Figure.1). All mutated structures were aligned and overlapped with the crystal structure to create the RMSD plot. The reported RMSD value of katG is 0.031Å. The crystal structure of pncA protein nicotinamidase/pyrazinamidase was obtained from RCSB PDB server corresponding to PDB id 3PL1 (Figure. 2). Two mutant structures were generated from 3PL1. The 3PL1 structure was aligned and overlapped with the two mutant structures (Figure.2). The recorded RMSD value of pncA was 0.082Å. rpoB protein DNA-directed RNA polymerase β-subunit was obtained from the RCSB PDB server corresponding to PDB ID 5UHC (Figure.3). Five mutant structures were generated corresponding to the 5UHC in respective positions. These obtained mutant structures were aligned and overlapped with the crystal structure 5UHC (Figure.3). Pair wise RMSD plot was created with 0.014Å in MOE. DNA gyrase subunit A protein is encoded by gyrA gene. The mutated protein structure of crystal structure 3IFZ was created in protein builder (Figure.4). The crystal and the mutant structure were aligned and pair wise RMSD plot was created in MOE. Overlapping structures reported 0.002Å RMSD. Different mutations in genes could be linked to multidrug resistance.

**Figure. 1:**
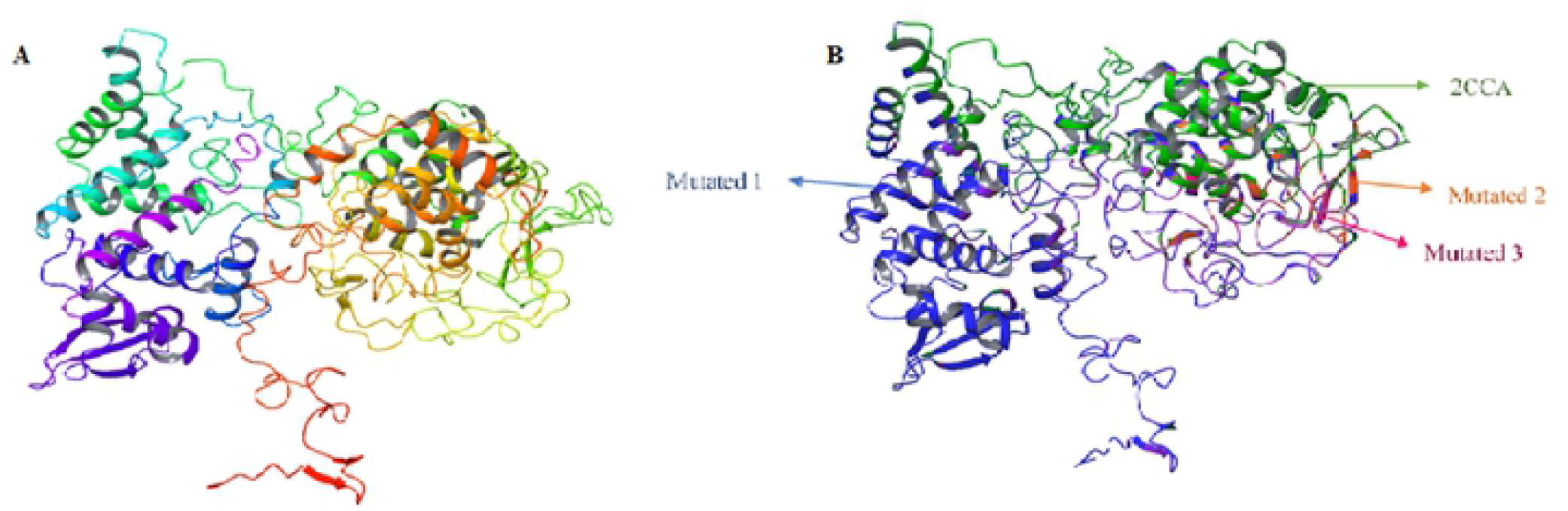
[A] Crystal structure of 2CCA. [B] superpose structures of 2CCA and mutated structures

**Figure. 2:**
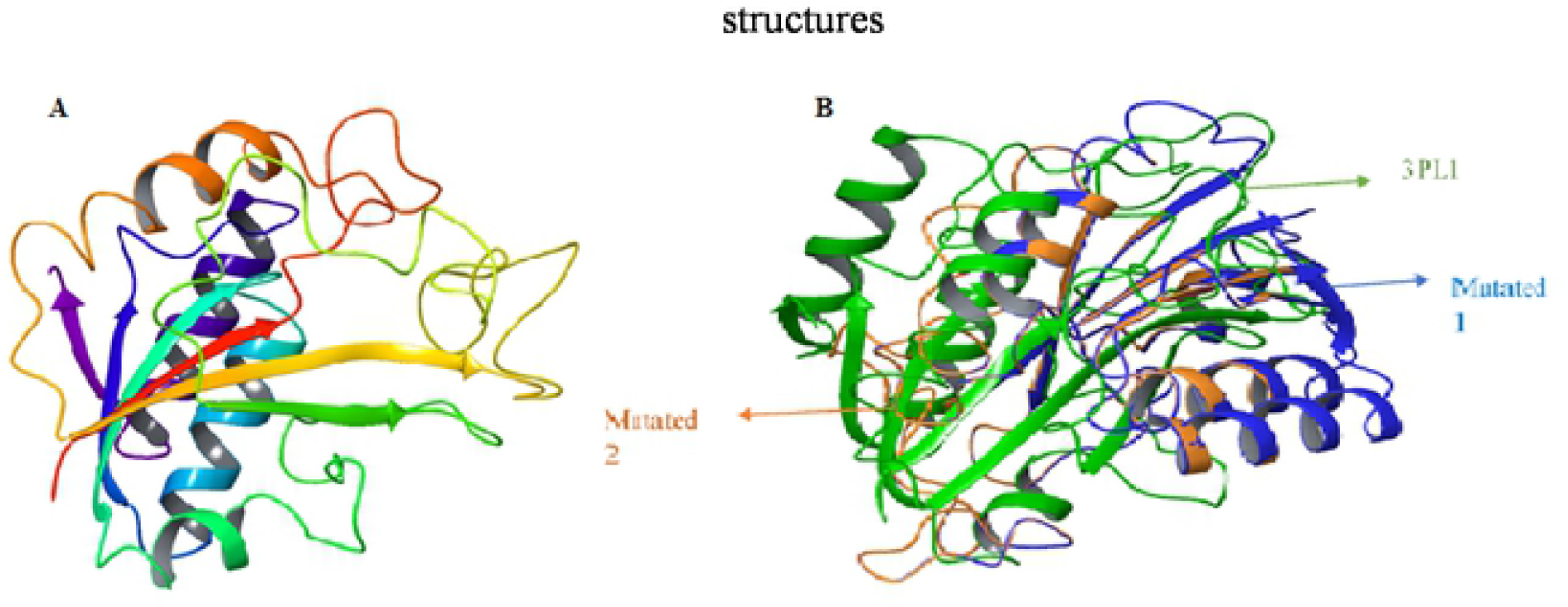
[A] Crystal structure of 3PL1. [B] Superpose structure of 3PL1 with mutated structures

**Figure. 3:**
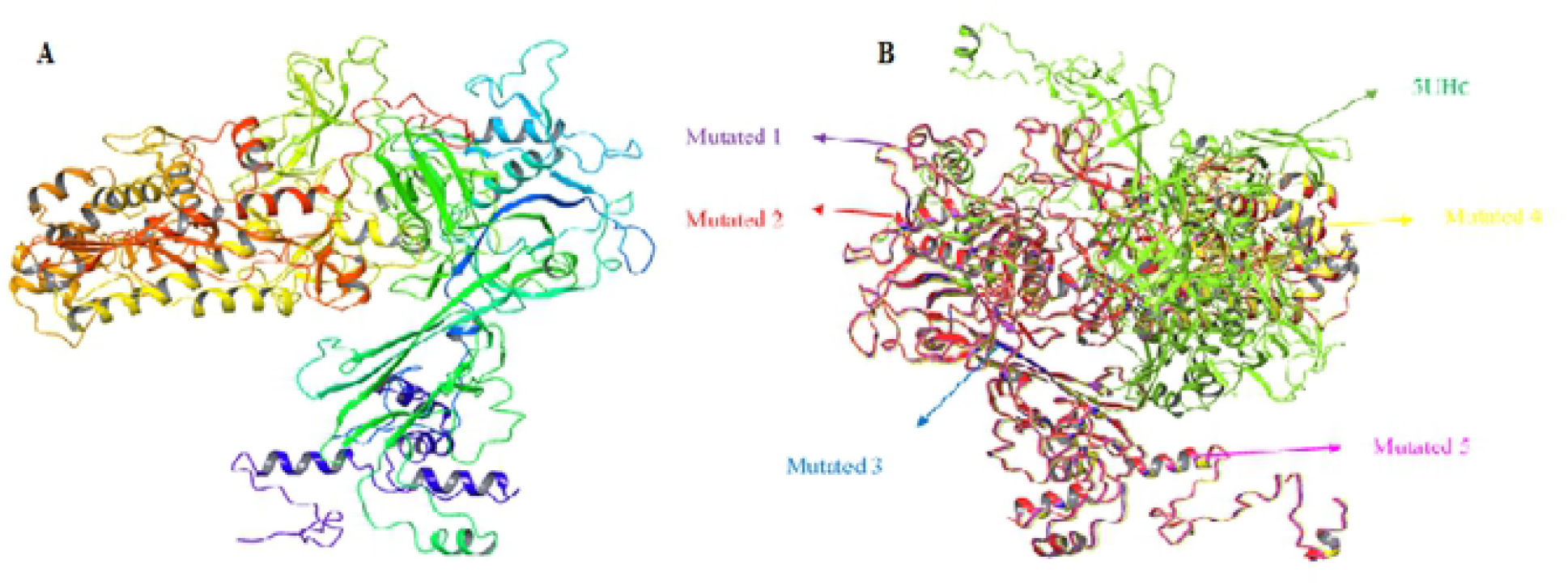
[A] Crystal structure of 5UHC [B] Superposed structure of 5UHC with mutated 5UHC

**Figure. 4:**
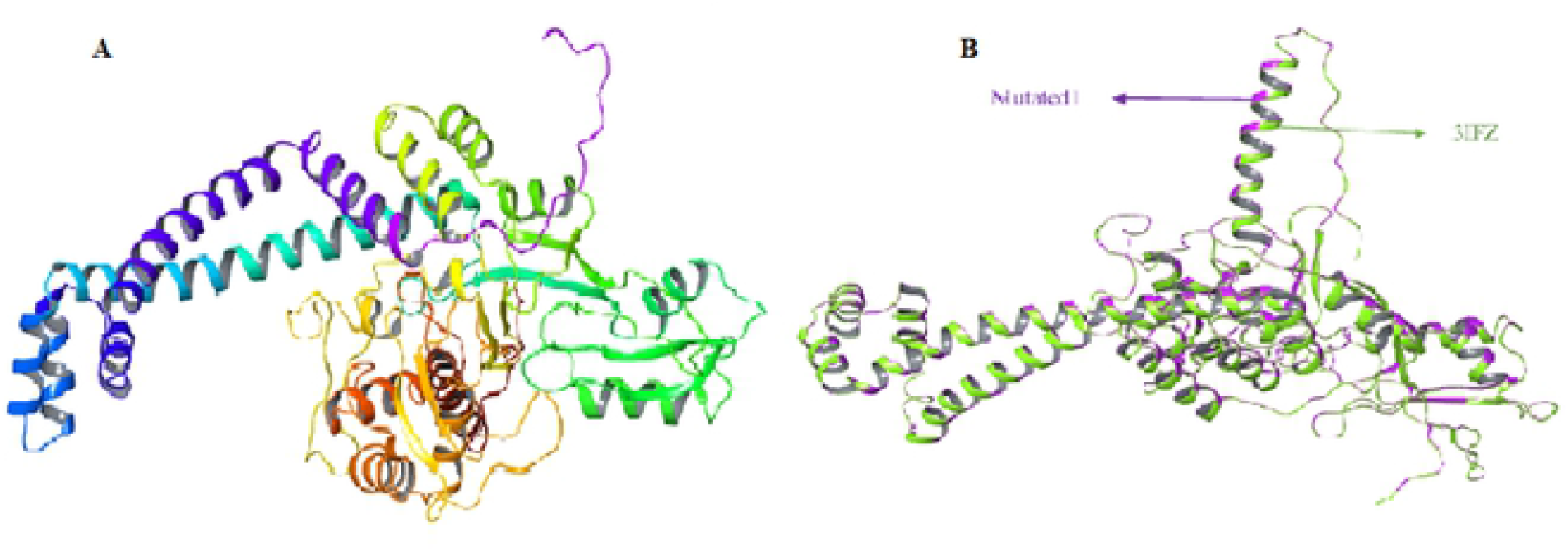
[A] Crystal structure of 3IFZ [B] 3IFZ crystal structure aligned and super posed with mutated 3IFZ

### Pharmacophore Generation

Isoniazid is one of the reported ligands for the katG protein. The 3D structure of the ligand was acquired from the PubChem database corresponding to CID number 3767. The obtained ligand was protonated and reduced in MOE with an MMFF94x force field. The pharmacophore was created in MOE. Figure-5 shows the pharmacophoric characters of isoniazid. For isoniazid, three features are hydrogen bond acceptor and two features of hydrogen bond donor, one hydrophobic interaction and the benzene ring interaction was also selected as aromatic interaction. The reported ligand for pncA protein is pyrazinamide. The 3D structure of the pyrazinamide was obtained from PubChem database corresponding to CID number 1046. The ligand was minimized and protonated in MOE and the pharmacophoric properties were constructed on the structure of pyrazinamide. The pharmacophoric properties of pyrazinamide are shown in Figure– 5. The pyrazinamide pharmacophore has 3 hydrogen bond acceptor features and 1 hydrogen bond donor feature, two hydrophobic interactions and benzene ring was selected as an aromatic interaction feature. The reported ligand for rpoB protein is rifadin. The 3D structure of rifadin was obtained from the PubChem database corresponding to CID number 5381226. The ligand was minimized and protonated and pharmacophoric properties were created on rifadin in MOE. Figure–5 presents the pharmacophoric properties of Rifadin. The pharmacophoric features of rifadin include three hydrogen bond acceptor features, one hydrophobic and one aromatic interaction. The DNA gyrase subunit A already carries a reported ligand: ofloxacin. The 3D structure of the ofloxacin was obtained from the PubChem database corresponding to CID number 4583. The ligand was minimized and protonated and pharmacophoric features were created on ofloxacin structure in MOE. Figure–5 represents the pharmacophoric properties of ofloxacin. Three features were selected as hydrogen bond acceptor, one hydrophobic interaction, one hydrogen bond donor and 2 aromatic interactions were generated with MOE2016.

**Figure 5:**
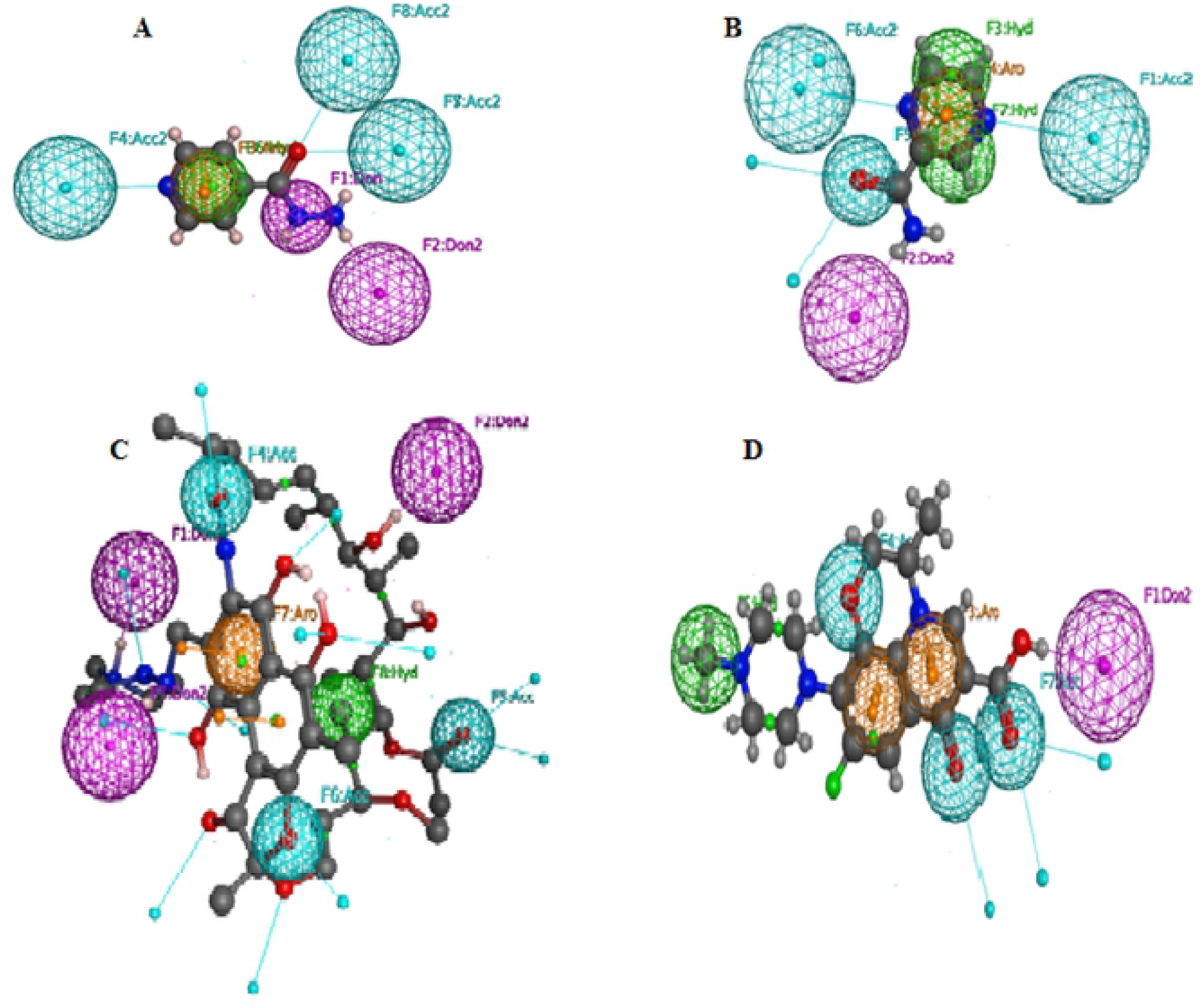
[A] Pharmacophoric features of isoniazid [B] Pharmacophoric features of pyrazinamide [C] Pharmacophoric feature of rifadin [D] Pharmacophoric features of ofloxacin

### Complexed base Pharmacophore Screening

The identified pharmacophoric features of all the 4 ligands were applied for screening the database in order to find small molecules with equivalent properties. All pharmacophoric features were set as essential for screening the ZINC small molecule drug like database, which contains 17,900,742 small compounds, was chosen for pharmacophore base scan. Small molecules, which qualify the set standards of the pharmacophore model, were elected for a new database. Total hits of 486 small compounds for katG ligand pharmacophore were obtained from ZINC database screening. The pncA protein ligand pharmacophore resulted in 542 hits of small drug like compounds from ZINC database screening. The rpoB protein ligand pharmacophore screening provided 112 hits of small compounds from the ZINC database. The gyrA protein ligand pharmacophore screening resulted in 365 hits for small compounds from ZINC database. Pharmacophore screening is a very useful and suitable tool for new drug discovery that enables finding potent and novel compounds.

### Molecular Docking

Molecular docking increases the probability of appropriate hits of molecules. Prior to docking the output databases into the protein active sites, all the four output databases were subjected to Lipinski’s rule. All screening outputs from ZINC database were evaluated by London dG scoring method in MOE2016 and resulting five conformations were selected for each ligand using the proxy triangle algorithm. Of these docked conformers 15% were selected on the basis of docking score, which was further analyzed for binding interactions. Finally, on the basis of docking score and binding interactions, ten small molecules were selected for each protein as novel inhibitors. Three-dimensional representation of the interactions of compounds with active pocket of proteins is shown in Figure-6, 7, 8 and 9.

**Figure 6:**
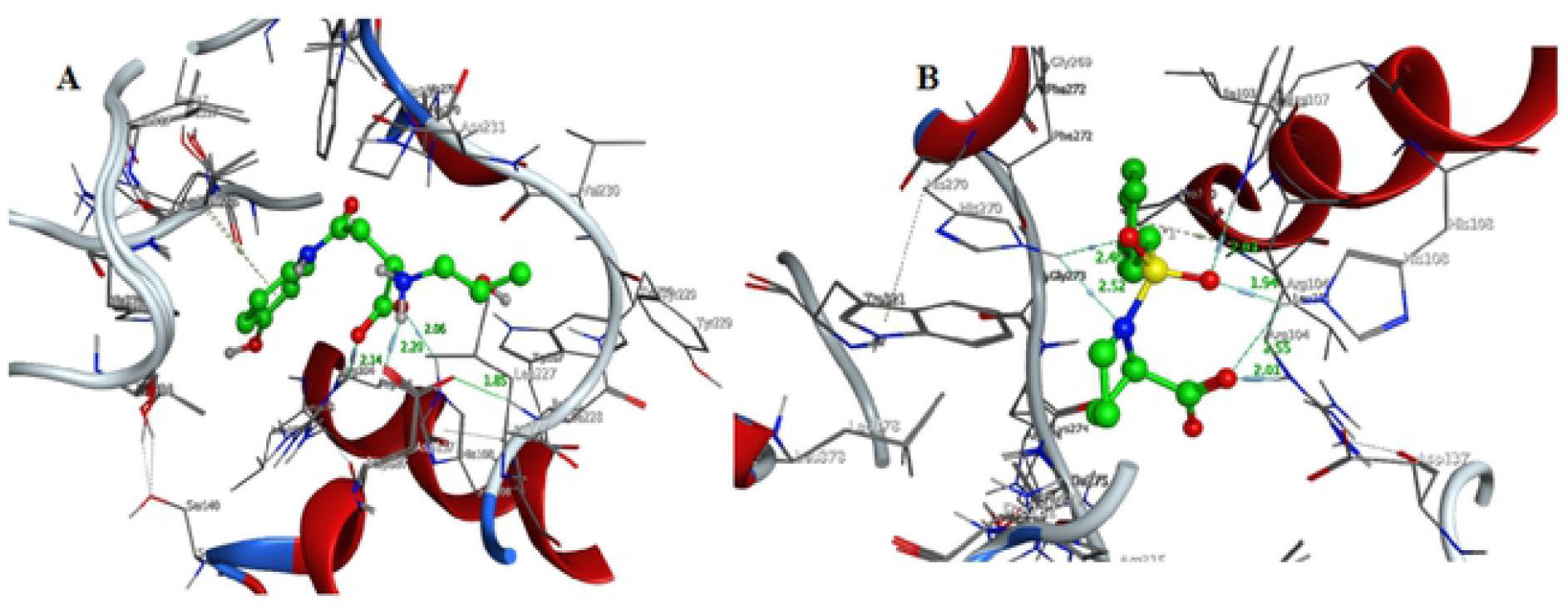
Three-dimensional representation of the interactions of compound ZINC05257859 (A) and ZINC11891015 (B) with katG active pocket.

**Figure 7:**
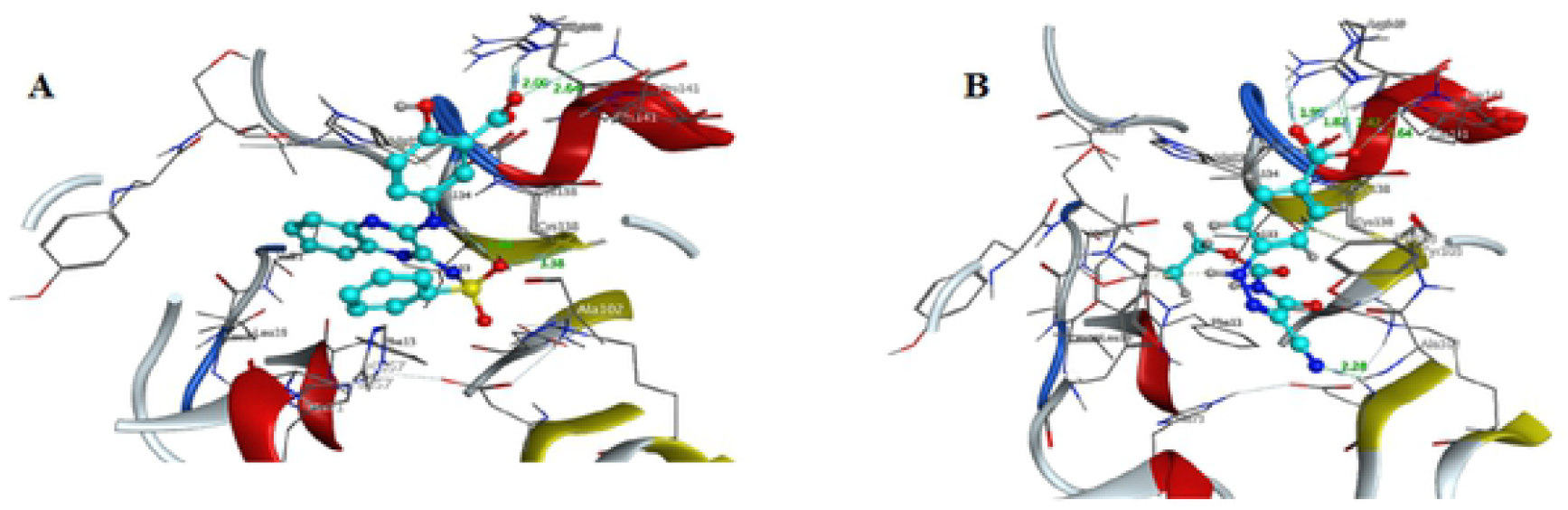
Three-dimensional representation of the interactions of compound ZINC20828864 (A) and ZINC05459404 (B) with pncA active pocket.

**Figure 8:**
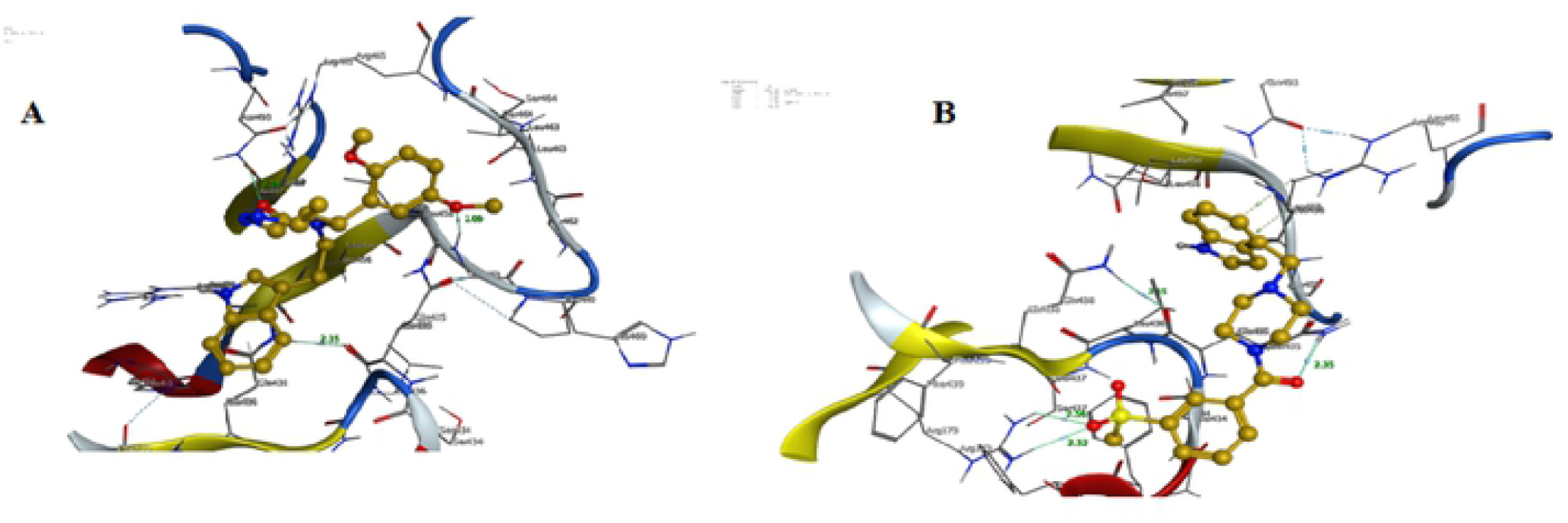
Three-dimensional representation of the interactions of compound ZINC39272743 (A) and ZINC93480289 (B) *with rpoB* active pocket

**Figure 9:**
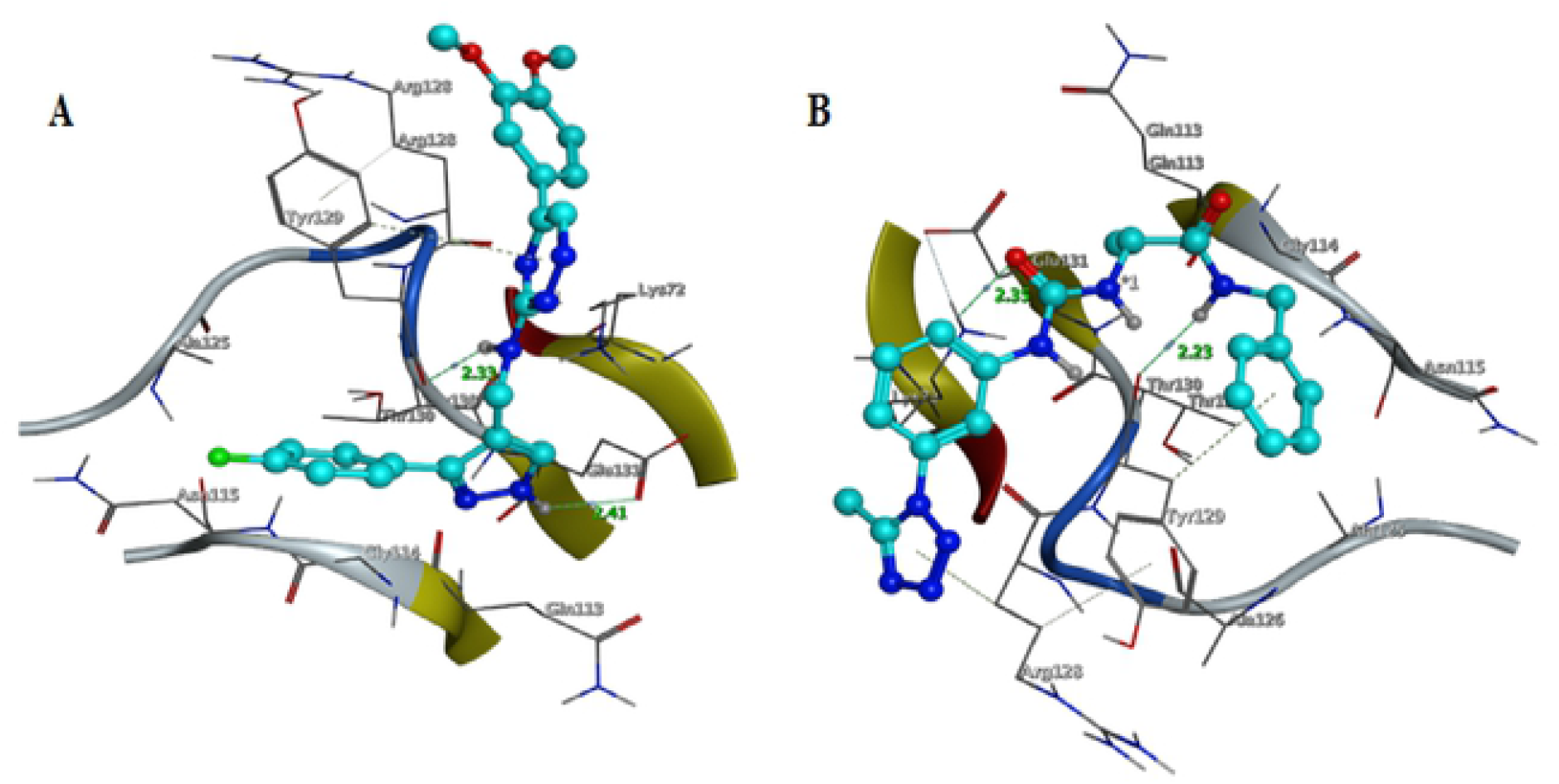
Three-dimensional representation of the interactions of compound ZINC12433306 (A) and ZINCS6640502 (B) with gyrA active pocket.

### Docking Score and Binding Energies

Binding energies and binding affinities of all the four proteins and ligands, which were obtained from ZINC databases docking score, were calculated by the MOE software and are reported in kcal/mol in Table–1-4.

**Table 1:** ***katG*** final hits docking score and RMSD refine

| **S No** | **Name** | **Docking Score** | **RMSD Refine** | **E Score** |
| --- | --- | --- | --- | --- |
| **1** | ZINC88613615 | -129.4315 | 1.6020 | -17.3896 |
| **2** | ZINC73694963 | -111.2725 | 1.2888 | -15.2426 |
| **3** | ZINC13564131 | -107.2721 | 1.9801 | -22.0461 |
| **4** | ZINC48331854 | -105.8490 | 1.9397 | -16.5574 |
| **5** | ZINC92463385 | -105.1475 | 1.1592 | -12.3351 |
| **6** | ZINC92479119 | -101.8164 | 1.7324 | -14.2246 |
| **7** | ZINC00205908 | -91.6005 | 1.1732 | -14.7671 |
| **8** | ZINC05257859 | -91.2623 | 1.7630 | -21.1987 |
| **9** | ZINC35303163 | -88.5217 | 1.9898 | -15.0075 |
| **10** | ZINC11891015 | -83.6541 | 1.6294 | -14.1690 |

**Table 2:** *pncA* final hits docking score and RMSD refine

| **S No** | **Name** | **Docking Score** | **RMSD Refine** | **E Score** |
| --- | --- | --- | --- | --- |
| **1** | ZINC38290758 | -91.8779 | 1.3383 | -13.4888 |
| **2** | ZINC03844927 | -83.7522 | 1.1612 | -15.9408 |
| **3** | ZINC38290757 | -83.7206 | 1.9661 | -14.0429 |
| **4** | ZINC03844926 | -83.5078 | 1.4821 | -14.6584 |
| **5** | ZINC32095180 | -83.2028 | 1.2798 | -14.7901 |
| **6** | ZINC13409428 | -80.3618 | 1.4415 | -14.7268 |
| **7** | ZINC49474085 | -79.1390 | 1.5748 | -13.0041 |
| **8** | ZINC20828864 | -76.9627 | 1.6320 | -10.7699 |
| **9** | ZINC05459404 | -76.6925 | 0.7688 | -12.1186 |
| **10** | ZINC00392982 | -76.3667 | 1.3575 | -12.5163 |

**Table 3:** *rpoB* final hits docking score and RMSD refine

| **S No** | **Name** | **Docking Score** | **RMSD Refine** | **E Score** |
| --- | --- | --- | --- | --- |
| **1** | ZINC16927899 | -761.3942 | 1.6372 | -36.5957 |
| **2** | ZINC40475355 | -701.1836 | 2.0014 | -35.2079 |
| **3** | ZINC16735142 | -691.0308 | 0.9966 | -35.8693 |
| **4** | ZINC95486285 | -657.1195 | 1.2217 | -30.9879 |
| **5** | ZINC85412686 | -572.2875 | 1.3451 | -28.8134 |
| **6** | ZINC40475335 | -556.6644 | 2.7125 | -40.8213 |
| **7** | ZINC19762653 | -514.9138 | 1.8795 | -17.5004 |
| **8** | ZINC15657726 | -491.0989 | 1.1917 | -20.4539 |
| **9** | ZINC39272743 | - 482.3535 | 2.1498 | -16.1519 |
| **10** | ZINC93480289 | -446.7381 | 4.0505 | -18.1358 |

**Table 4:** *gyrA* final hits docking score and RMSD refine

| **S No** | **Name** | **Docking Score** | **RMSD Refine** | **E Score** |
| --- | --- | --- | --- | --- |
| **1** | ZINC72086818 | -53.7750 | 4.2784 | -32.1013 |
| **2** | ZINC71892626 | -53.7117 | 1.9045 | -28.9632 |
| **3** | ZINC11819004 | -52.6137 | 1.6921 | - 28.3720 |
| **4** | ZINC89928330 | -51.5949 | 1.7039 | -28.4039 |
| **5** | ZINC16955083 | - 51.530 | 0.9361 | - 29.1497 |
| **6** | ZINC09576286 | -51.4883 | 2.4492 | -31.3807 |
| **7** | ZINC13566986 | -49.4356 | 2.1497 | -31.3699 |
| **8** | ZINC13164785 | -48.4203 | 1.5245 | - 29.6528 |
| **9** | ZINC12433306 | -47.4112 | 1.2365 | -26.1476 |
| **10** | ZINCS6640502 | -44.3937 | 3.1432 | -26.5130 |

## Discussion

In this study, proteins encoded by katG, pncA, rpoB and gyrA gene of M. tuberculosis were selected and targeted by drug like small compounds from ZINC drug like databases using an in-silico approach. The data analysis generated numerous novel and potent small molecules against the proteins of katG, pncA, rpoB and gyrA gene.

Virtual screening based on the structure and ligand is an important tool in medicinal chemistry, which plays a significant role in identification and chemo informatics. These virtual screening methods are found to be widely applied to many therapeutic targets.

katG codes for the enzyme catalase/peroxidase, which is involved in activation of isoniazid. The reduced or absent activity of this enzyme occurs due to the mutations in katG gene, which is responsible for isoniazid resistance (Zhang et al., 1992). Pyrazinamide is a structural analogue of nicotinamide. It is converted to active form of pyrazinamide i.e. pyrazinoic acid by the enzyme pyrazinamidase that is encoded by pncA gene. Mutations in pncA gene contribute to pyrazinamide resistance in M. Tuberculosis. Rifampicin works by binding with RNA polymerase β-subunit encoded by rpoBgene and thus blocking the elongation of messenger RNA (Blanchard, 1996). DNA gyrase is an enzyme involved in the super coiling of DNA and consists of subunit A and subunit B. Ofloxacin, a synthetic antibiotic of the fluoroquinolone drug class, is the second line drug that works by binding to gyrase-DNA complexes and inhibiting DNA replication. Ofloxacin resistant strains develop through mutations in quinolone resistance-determining regions of gyrA and gyrB genes (Maruri et al., 2012). Using in silco approach about top 10 compounds were selected as potent and novel inhibitors on the basis of docking score and interaction for each protein. A total of 40 compounds, 10 for each protein from ZINC database, were chosen as promising lead compounds, which may act as novel, powerful, and structurally diverse inhibitors for four drug-resistant mutant proteins of M. tuberculosis. One of the related study was performed, targeting inhA and kasA genes and novel inhibitor was designed against the proteins responsible for mycolic acid synthesis present in M. tuberculosis (Sharma et al., 2015). Another related study was performed to discover novel inhibitors of M. tuberculosis inhA enzyme using in silico approach (Pauli et al., 2013). The present study reveals that the pharmacophore-based approach of screening base ligand can be convenient in the discovery of structurally diverse hits. These hits were structurally diverse with significant docking score and good binding affinity with the targeted proteins’ 3D structures.

The purpose of this study was to obtain a ligand having superior characteristics to inhibit the drug-resistant protein. The inhibitors generated as an outcome of molecular docking studies of pharmacophore screening allowed predicting new, potent and structurally different inhibitors for the mutant drug-resistant proteins of M. tuberculosis.

## Conclusion

Forty small compounds, ten for each protein were selected as final hits. The docking score of each novel inhibitor was found higher. The result indicates that novel inhibitors could be a promising lead compound and be effective in treating sensitive as well as multi drug resistant tuberculosis. We believe that these new scaffolds might be the good starting point for potential novel drugs and surely help the experimental designing of the anti-tuberculosis drug in a short time.

## Acknowledgments

The authors are grateful to Professor Abdul Wadood (Department of Biochemistry, Abdul Wali Khan University Mardan, Pakistan) for his useful suggestions and support during manuscript preparation.

